# High-quality transcriptome profile of *Treponema pallidum* subsp. *pallidum*: confirmation of transcriptional landscape

**DOI:** 10.1101/2024.08.22.609093

**Authors:** Linda Grillová, Eli M Carrami, William Roberts-Sengier, Nicholas R. Thomson

## Abstract

Syphilis remains a critical global health challenge due to its potential for severe complications and the increase in its incidence rate over recent years. Until recently, the infectious agent of syphilis, *Treponema pallidum* subsp. *pallidum* (TPA), could not be cultured *in vitro*. Advances in co-culture techniques have finally allowed for effective long-term cultivation of TPA, providing a platform to study its biology. Limited transcriptional data from TPA have been reported so far and many genes in treponemal genomes are annotated based on *in silico* prediction of putative coding sequences without functional validation. To inform future syphilis vaccine development, experimental validation of *in silico* predicted genes coupled with functional annotation is necessary. In this study, strand-specific RNA-sequencing was used to reconstruct a high-quality transcriptome profile of TPA, confirming the active transcription of genes previously annotated as hypothetical, paving the way for more accurate identification of vaccine target candidates. Our transcriptomic data also revealed, for the first time, the organization of genes into transcription units, an abundance of anti-sense RNAs, and transcripts from intergenic regions, providing crucial insights for future functional genomics studies of TPA.

**Author Summary:** In our study, we explored the genetic activity of the bacteria responsible for syphilis, *Treponema pallidum* subsp. *pallidum* (TPA). Although syphilis has been a known disease for centuries, the bacterium causing it has remained difficult to study because it couldn’t be easily grown in the lab. Recently, new techniques have allowed us to cultivate TPA successfully, enabling deeper investigation into its genetics. By employing directional RNA sequencing, we have mapped out which genes are actively transcribed, including those previously labeled as hypothetical. Our study has also revealed new insights into the gene organization and uncovered the presence of antisense RNA, which may regulate gene expression. These findings offer critical information that could inform future research and vaccine development efforts for syphilis.

## Introduction

Syphilis, caused by the spirochete *Treponema pallidum* subsp. *pallidum* (TPA), is one the oldest known diseases that can lead to irreversible neurological and cardiovascular complications, stillbirths and miscarriages. Despite the availability of effective treatments, the incidence of syphilis has increased substantially over the last decade and thus this historical disease remains a serious challenge for global health today [1, 2]. Without an effective vaccine available, syphilis prevention strategies are limited to safe sexual behaviours and intensive screening.

TPA has a small minimal genome that lacks multiple metabolic and biosynthetic pathways which are necessary for the *de novo* biosynthesis of various enzyme co-factors, fatty acids, and nucleotides, which TPA must acquire from the host [3]. As a result, difficulties in culturing this pathogen *in vitro* and the low levels of TPA loads in clinical samples have limited our understanding of syphilis biology and pathogenesis. Several years ago, after decades of experiments with various co-culture systems [4], in a ground breaking discovery, Edmondson and colleagues reported the first successful long-term cultivation of TPA co-cultured with rabbit skin epithelial (Sf1Ep) cells [5]. This co-culture system has opened new avenues for the study of the basic biology of pathogenic treponemes and has allowed to gain new knowledge on the susceptibility/resistance of TPA to clinically relevant antibiotics [6-9], differential gene expression in TPA strains cultured *in vivo* versus *in vitro* [10], and the mechanism behind infection persistence in absence of treatment [11]. Furthermore, the recently introduce methodology for genetic engineering of TPA opened new opportunities for the evaluation of gene functions in these elusive bacteria [12].

To inform syphilis vaccine development, experimental validation of *in silico* predicted genes coupled with functional annotation is necessary. Currently, however, many TPA genes are annotated based on *in silico* prediction of putative coding sequences (CDSs) without a corresponding functional validation to demonstrate that these CDSs are indeed transcribed and expressed. Therefore, to guide future functional genomics studies of TPA, in this study we aimed to reconstruct a high-quality transcriptome profile of TPA using strand-specific RNA-sequencing. These data allowed us to elucidate the transcriptional landscape of TPA and provide evidence that most genes are transcribed, including those annotated as hypothetical proteins. In addition, we identified, for the first time, the organization of genes into transcriptional units, an abundance of anti-sense RNAs (asRNA), and various transcripts arising from intergenic regions (IGR).

## Results

The main goal of this study was to obtain a high-quality data to profile TPA transcriptome and correlate it to the current genomic annotation for this pathogen. Therefore, we first performed whole-genome sequencing of the TPA strain Nichols cultured in our laboratory to identify nucleotide variants relative to the genomic reference available for this strain. Subsequently, we sequenced the Nichols’ strain transcriptome to a high depth of coverage and used that to validate the published genome annotation (NC_021490.2).

### Whole genome sequencing and preparation of a custom reference genome

To generate a highly accurate reference genome for our test strain, the SureSelect Target enrichment system was utilized to enrich for TPA-specific DNA used to construct the sequencing libraries, as described previously [13]. Over 39% of the total reads, amounting to 4.5 million, mapped to the genomic reference NC_021490.2, resulting in an average depth coverage of 555x. Only base variants with frequencies exceeding 50% were considered. In total, 15 single nucleotide variations (SNVs) were identified when comparing the genomic sequence of the *in vitro* cultured Nichols to the archetypal reference genome NC_021490.2. The majority of SNVs were located in coding regions (n = 11), with 7 predicted to lead to non-synonymous amino acid substitutions. Additionally, we detected a 67-nt deletion in the intergenic region upstream of *tp0895* gene, which appears to have no functional consequences as it does not affect the transcription of neighboring genes, and a 126-nt insertion in the gene encoding the fibronectin-binding protein (TP0136), resulting in a frameshift and premature truncation of the predicted protein [14]. All these variations were described in previously sequenced strains belonging to the Nichols clade (e.g., Nichols Seattle – CP010422 and Dallas) [15]. Heterogeneity was observed in several poly-G and poly-C homopolymeric tracks, mostly occurring in predicted promotor regions, also previously reported [16]. Two changes in the lengths of these homopolymeric tracts, however, were detected in coding regions, both predicted to cause frameshifts in genes predicted to encode proteins of unknown function (S1 Table).

The custom reference genome for the subsequent analyses was constructed by transferring all annotations from the RefSeq NC_021490.2 (updated as of 03/2023) and incorporating the aforementioned nucleotide variations and the resulting changes into the annotation. In line with generally adopted coding conventions, we used locus tag IDs as originally described in the reference genome CP004010.2 [17] wherever possible. The new custom reference genome had a length of 1,139,617 bp and contained 1041 annotated features including 987 coding DNA sequences (CDSs), and 54 RNAs (including 6 rRNAs, 45 tRNAs, 2 ncRNAs and 1 tmRNA).

### Directional RNA libraries obtained from in vitro cultured TPA

To reveal a higher breadth of transcripts expressed in TPA cells, we extracted RNA from TPA cultivated both under standard *in vitro* co-culture conditions and conditions designed to stress TPA cell growth: incubated in media without the presence of mammalian cells (“axenic culture”). Since the presence of mammalian cells is required for the continual growth of TPA, as expected, the number and the motility of TPA cells after a week of cultivation differed between these two conditions (see details in Materials and Methods). However, because TPA is known to remain metabolically active in the absence of mammalian cells for up to seven days [5], the quality of the extracted RNA was high in both conditions (RIN>8).

For library preparation, the “co-culture” RNA sample was ribo-depleted using probes against both the eukaryotic and the prokaryotic rRNAs to reduce the amount of rRNA transcripts. Additionally, this RNA sample was also used for poly(A)-depletion using oligo-dT magnetic beads, to reduce the contribution of eukaryotic mRNAs and further enrich TPA mRNAs in the sample. To account for any TPA transcripts that may have been compromised due to poly(A)-or prokaryotic ribo-depletion, we treated a second aliquot of the “co-culture” RNA sample with only the eukaryotic ribo-depletion probes. The “axenic culture” RNA sample was only used for prokaryotic ribo-depletion. Subsequently, three directional RNA libraries were prepared using (I) ribo- and poly(A)-depleted “co-culture” RNA, (II) only ribo-depleted “co-culture” RNA and (III) ribo-depleted “axenic culture” RNA (Fig 1). All libraries were of a high quality and were deep sequenced using over 100 million reads for each library to increase the representation of low-abundance transcripts.

**Fig 1.**
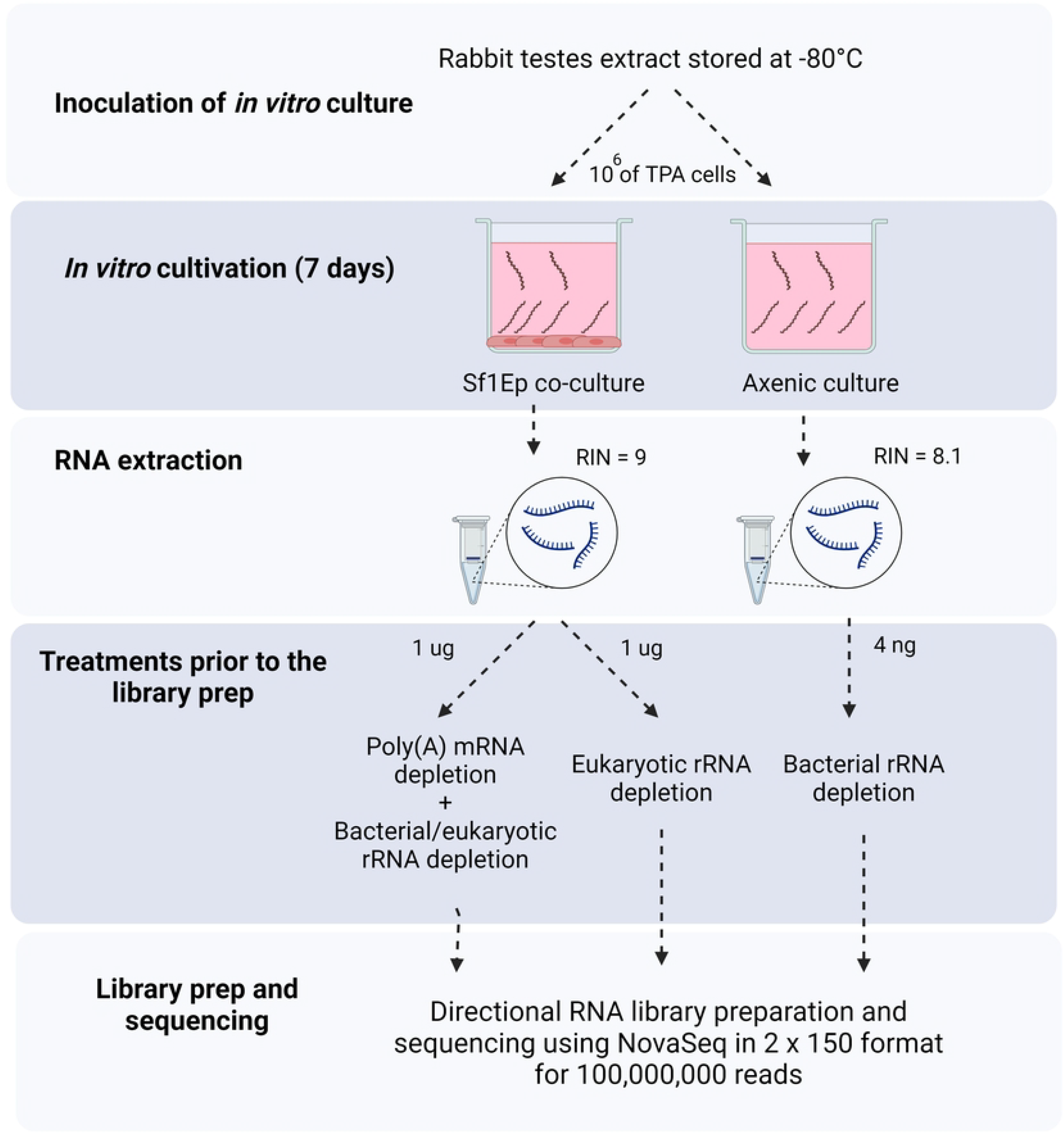
The workflow for strand-specific RNA sequencing of *in vitro* cultured TPA for deep transcriptome sequencing. One million bacterial cells from an *in vivo* culture were inoculated in TPCM-2 media both in the presence or absence of Sf1Ep cells, and were incubated in low oxygen incubator (1.5% of oxygen). After 7 days, the RNA from both samples was extracted, and prior to library preparation, were treated as indicated. Multiple different treatments were performed before library preparation to ensure the transcripts would not be compromised. Directional RNA library preparation was performed, and each library was sequenced using Illumina NovaSeq to generate 30 Gbp of data per sample (100 million PE150 reads). Created with BioRender.com.

For transcriptome reconstruction, the reads from all three libraries were first merged and mapped to the new custom reference. Aligned reads were then de-duplicated resulting in over 5 million unique paired-end reads covering over 99% of the TPA genome. To gain an overview of the TPA expression profile, we temporarily masked paralogous regions of the TPA genome and plotted the coverage values across the rest of the genome (Fig 2A). Due to the long stretches of identical sequences in the paralogous regions, Illumina short reads cannot be accurately assigned to either member of the paralogous pair, thereby resulting in ambiguous mapping (Fig 2A, S2 Table). However, after excluding reads with multiple mappings from the analysis, transcription was confirmed in all paralogous regions by identifying read coverage in unique segments of these regions. These results supported that nearly all genes in the TPA genome are expressed as described in more detail below.

**Fig 2.**
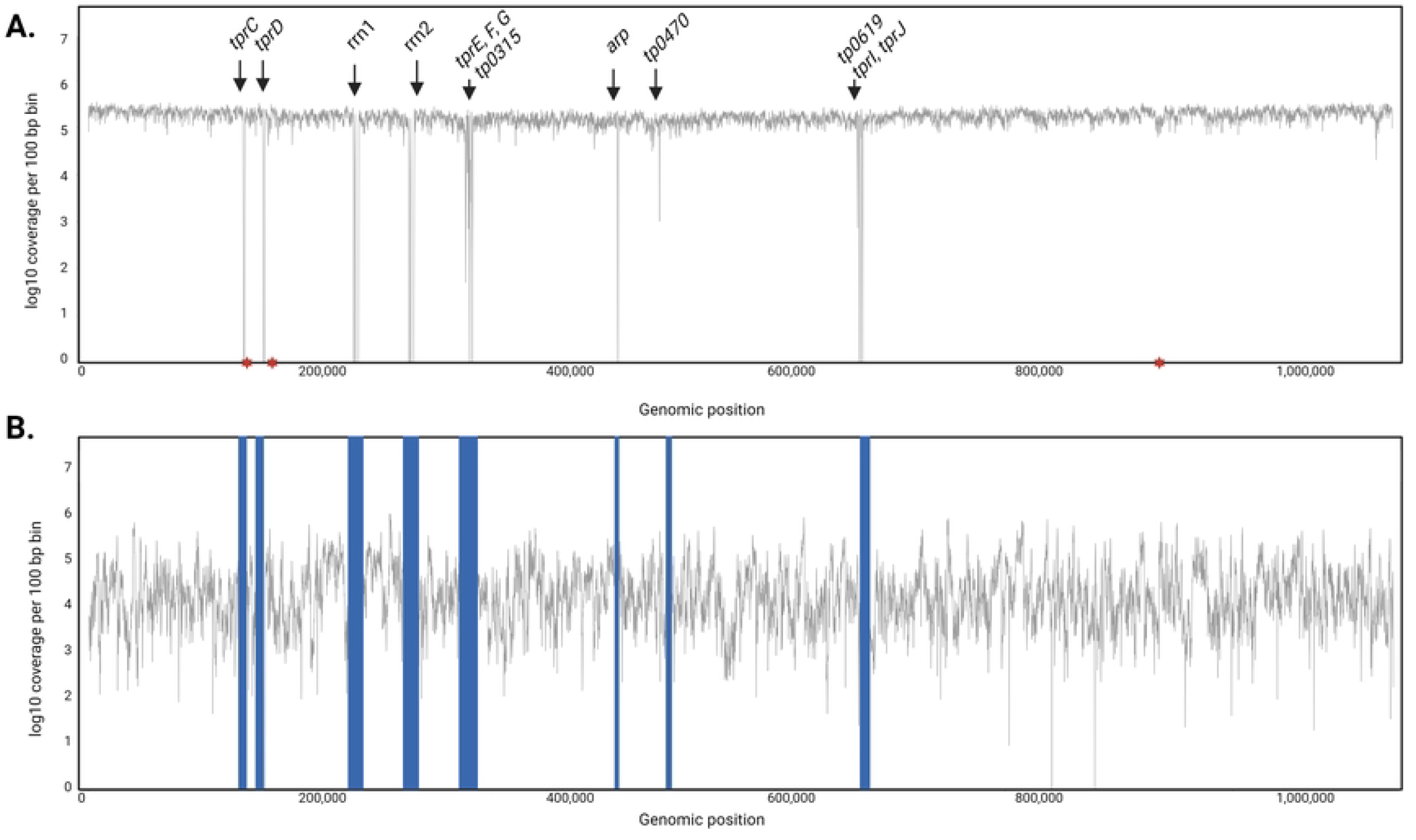
Log10 coverage per 100 bp across the TPA genome and transcriptome. **A**. Genome coverage with arrows highlighting long identical sequence stretches that lead to ambiguous mapping. The apparent zero coverage is because reads with multiple mappings were excluded from the analysis. Asterisks indicate the location of frameshifts leading to the modification of annotation. **B**. The transcription profile of the TP genome, with paralogous regions masked, displays a uniform pattern, suggesting stable transcription across most annotated regions of the TPA genome.

### Most annotated regions in TPA genome are transcribed

Further analysis of the RNA sequencing results showed that of the 1041 annotated features in the TPA genome, the transcription of 1019 (98%) was supported by the presence of directional Illumina short read data corresponding to the correct orientation of the annotation (at least 10 correctly oriented unique reads spread across the gene) (Fig 2B). This included 99% of the mRNAs (973/987), 82% of the tRNAs (37/45) and 100% of rRNAs, ncRNAs and the tmRNA. The 8 missing tRNA transcripts were found to be correspond to tRNA genes with sizes ranging from 72 to 74 bp which were likely excluded during the library processing steps due to their small size. Importantly, more than half of the gene annotations where we found no convincing evidence that they were transcribed in the expected orientation (57%, 8/14) were annotated as encoding hypothetical proteins, while the rest coded for proteins involved in various metabolic and biosynthetic pathways (S3 Table).

Despite the inconsistencies above this analysis also provided first evidence that most genes annotated as coding for hypothetical proteins (138/146, S4 Table) are in fact transcribed, the existence of some of which has never been experimentally proven before. Furthermore, our results provide strong evidence for the expression of genes that offer as potential target candidates for syphilis vaccine development (n = 50) (S5 Table). Additionally, all annotated pseudogenes were actively transcribed (n = 5) suggesting possible miss-annotation of the functional variants of these genes, that they may still lead to the translation of a protein that itself may be functional perhaps with a potential role in the genome regulation of TPA. To simplify the visualization of the strand-specific transcription tracks, we generated directional coverage plots for all annotated regions of the genome and all intergenic regions larger than 10 bp (Fig 3A). This resulted in 1775 tracks, accessible at https://tpa-coverage-plots.pam.sanger.ac.uk/ (read coverage details in the S6 Table).

**Fig 3.**
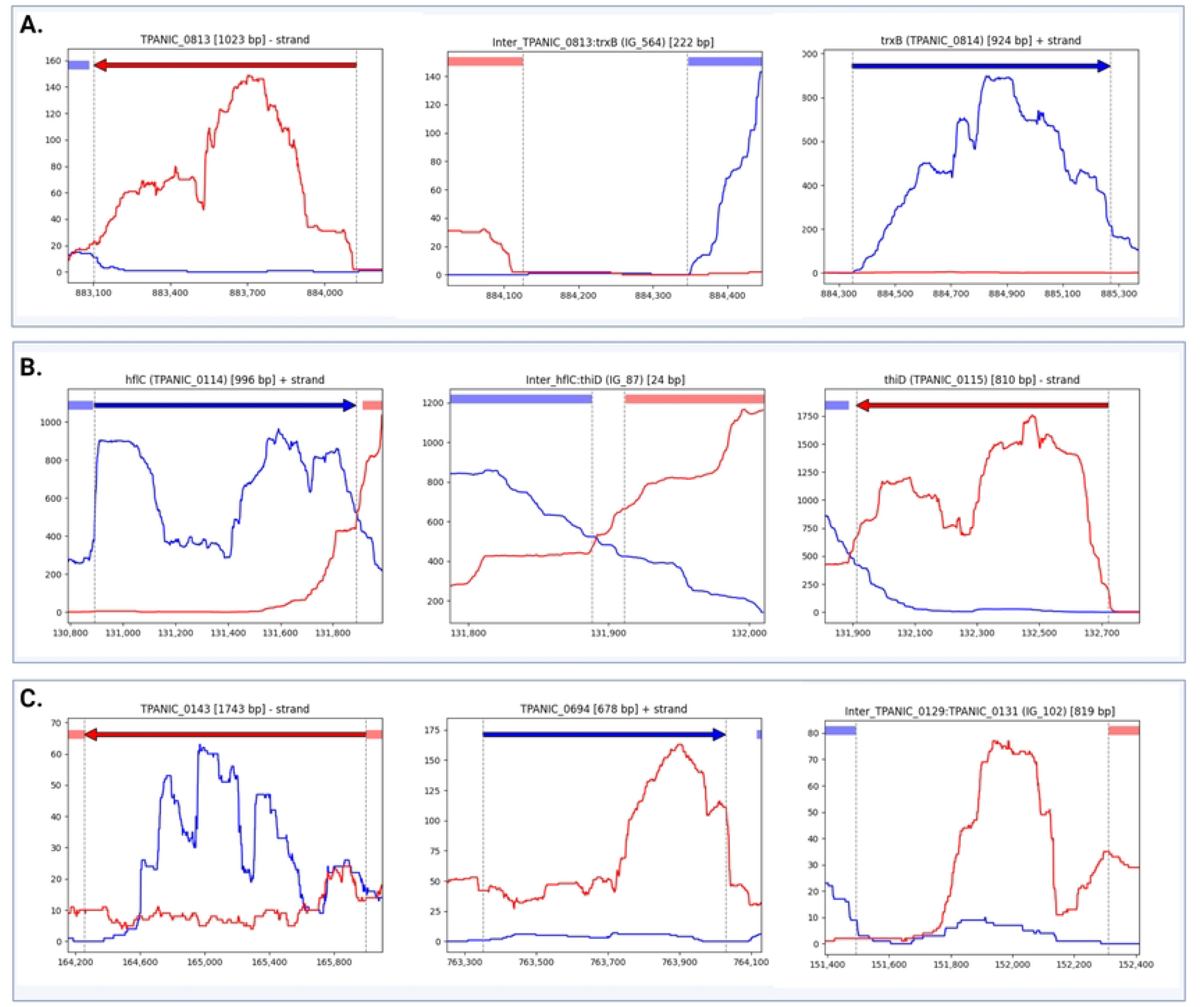
Strand-specific coverage plots. **A**. Blue and red traces show transcription in the plus and minus strand, respectively. Similarly, blue and red arrows on the top indicates the annotated genes in the plus and minus direction. The x-axes record the position on the genome and the y-axes record the number of reads mapped at the location (average coverage depth = 658). Highlighted regions on the top show neighbouring or/and overlapping genes. Examples of gene annotated and transcribed in the minus strand (*tp0813*, left), intergenic region (*tp0813*-*tp0814*, middle), and gene annotated and transcribed in the plus strand (*tp0814*, right). **B**. Example of a convergent 3’-3’ gene pair with overlapping transcripts. Transcription of the gene *tp0114* (left), intergenic region (*tp0114* - *tp0115*, middle) ad transcription of the gene *tp0115* (right). **C**. Example of transcripts that conflicted with existing annotations. Antisense transcript originating within annotated gene (*tp0143*, left), example of a region transcribed exclusively in the opposite orientation to the original annotated gene *tp0694* (middle), and example of transcripts arising from intergenic regions (*tp0129*-*tp0131*, right). The complete set of 1775 strand-specific coverage plots are accessible from https://tpa-coverage-plots.pam.sanger.ac.uk/ and read coverage details can be found in the S6 Table.

### Antisense transcripts and transcripts in conflict with the accepted annotation

Of the annotated regions, 43.7% (455/1041) were covered by both sense and antisense reads (mean coverage ≥ 5 reads). Moreover, 101 genes had an antisense:sense ratio of ≥ 0.5, and 72 genes had a ratio of ≥ 1. This means that over 7% of the genes in TPA genome are transcribed along with RNA transcripts from the antisense strand at similar or higher coverage levels as their sense strand. The visual inspection of strand-specific coverage plots indicated that most of the identified antisense transcripts were found to be arising from an oppositely oriented adjacent or overlapping gene (Fig 3B). Out of the total number of convergent 3’-3’gene pairs that can be found in the TPA genome (n = 125; with intergenic separation of 100 nucleotides or less), 107 genes (85%) contained antisense RNA arising from the oppositely oriented neighbouring genes.

In total, we identified 98 transcripts that conflicted with existing annotations. This included 79 antisense transcripts of varying lengths, mapping to positions within annotated genes and ranging from short sequences covering parts of the gene to longer sequences spanning multiple annotated genes (Fig 3C, S7 Table). Notably, 9 regions were transcribed exclusively in the opposite orientation to the original annotation (Fig 3C, S8 Table). Additionally, we found 10 transcripts arising from intergenic regions. Interestingly, more than half of the transcripts in conflict with existing annotations were found to contain open reading frames (ORFs) exceeding 250 nucleotides in length. Some of these ORFs matched annotations identified in different versions of Nichols genome annotation but are missing from NC_021490.2 (n = 4, S10 Table). These ORFs overlap with current annotations in NC_021490.2 and predominantly encode hypothetical proteins.

### Gene organization in transcriptional units

In addition to refining the annotations, ANNOgesic was used to predict gene organization based on transcriptional profiles and the prediction of DNA sequences that signal the end of transcription without requiring the Rho protein (Rho-independent terminator). These data were then manually verified, enabling the identification of 165 polycistronic mRNA transcripts, encompassing nearly 80% (n=822) of the genes annotated in the TPA genome. The transcriptional units contained variable numbers of annotated genes ranging from 2 to 41. Most of the transcriptional units (n=149, 90%) contained up to 10 genes. The average length size of these polycistronic mRNA molecules was 5.343 kbp (median = 3.428 kbp and ranged from 479 bp to 33.305 kbp). The complete list of operons and their genes is reported in S11 Table. As expected, the largest operons predicted to encode the most CDSs (41 and 31, respectively) carried genes encoding ribosomal proteins. Other long transcriptional units (n = 21) encompassed highly conserved gene clusters whose product are essential for cell division and cell wall biosynthesis functions as well as ribosome biogenesis and flagellar proteins.

## Discussion

The lack of extensive experimental validation for *in silico* gene predictions is still a near-universal problem in biology that influences all downstream analyses. This problem can be significantly circumvented by elucidating an organism’s transcriptome that, in turn, allows to identify with higher confidence the functional elements of the genome. We therefore reconstructed a high-quality transcriptome of TPA and correlated our data with the current genome annotation for this pathogen. Surprisingly, most of the annotated regions were found to be transcribed (98%), including the *in silico* predicted genes coding for hypothetical or uncharacterized proteins taxonomically restricted to *Treponema* (S4 Table). In addition, we have confirmed the transcription and transcriptional directionality for potential candidates for a syphilis vaccine development. These vaccine candidates were selected based on previous *in silico* and experimental observations indicating their potential as outer membrane proteins (OMPs) (S5 Table). Overall, our findings suggests that the TPA genome encodes nearly minimal set of genes that are all transcribed and likely essential for the bacteria to survive in the host or in culture. This is complementary to the study published by Šmajs and colleagues in 2005, revealing the transcriptome-wide pictures of mRNA molecules of TPA using probe-dependent cDNA sequencing [18]. However, compared to the previous study, the use of strand-specific RNA-seq allowed to define the directionality of transcripts and to identify transcripts that were not present in the original annotation. Some of these transcripts corresponded to the proteins previously annotated in different versions of Nichols genome annotation (S10 Table). For example, the negatively oriented *tp0451* gene annotated in the NC_021490.2 as hypothetical protein was not transcribed in our condition. Instead, we identified transcription from the opposite strand that corresponded to the *tp0451a* annotated in CP004010.2 as nucleobase cation symporter-2.

Interestingly, all annotated pseudogenes were actively transcribed (n = 5), suggesting potential misannotation of functional gene variants. Pseudogene annotation in the RefSeq annotation file, which is periodically updated by the NCBI through an automated pipeline approximately twice a year, varies with each version. Alternatively, these pseudogenes might produce RNA transcripts that regulate their respective parental genes, as previously described in other [19, 20]. Additionally, some pseudogenes with disrupted mRNA could be successfully translated, as observed in *Salmonella enterica* [21]. In TPA, phenotype changes were noted in pseudogene knockout (*tpr*A) strains, which grew significantly faster *in vitro* compared to the wild-type strain [22]. Moreover, Houston and colleagues recently detected pseudogene expression in the proteome of *in vivo* grown TPA [23]. These observations, combined with data from this study, indicate that TPA pseudogenes may possess functional roles, albeit not coding for a functional protein. Therefore, it is necessary to reconsider their potential roles, whether as defunct genes, having new regulatory functions, or potentially interfering with cellular processes.

Importantly, we have identified an abundant presence of antisense RNA (asRNA) transcribed widely across the whole TPA genome. Most studies of antisense RNA (asRNA) in bacteria are relatively recent. However, antisense transcription has been identified in many microbial species, and the proportion of protein-coding genes with antisense transcription ranges from 0 to 35.8% across 47 species [24]. In TPA, almost 44% of the annotated regions were covered by both sense and antisense reads, which places TPA among the bacteria with highest abundance of asRNA. This should be taken into consideration when designing primers for reverse transcription-qPCR to quantify transcripts of interest. Even when stranded cDNA is created, qPCR can amplify both strands, potentially leading to misinterpretation of the data. Antisense transcripts in bacteria can regulate the expression of their target genes in different steps of gene expression. This includes transcription interference or attenuation, translation stimulation or inhibition, and RNA degradation [25]. In TPA transcriptome, many antisense transcripts were found to be arising from an oppositely oriented adjacent or overlapping gene. This tail to tail orientation seems to be the most common type of asRNA in bacteria and probably plays crucial role in the coordination of bacterial gene expression. It has been shown for example that this overlapping transcription can silence the expression of neighbouring genes or operons that code of opposing function [25, 26]. Other asRNA was found to be arising within the genes. While most of these transcripts likely represent RNA molecules with regulatory functions, some may represent transcripts coding for peptides or proteins that were not described before (misannotation and/or overlapping genes). In fact, more than half of the transcripts identified as conflicting with the current annotation could be matched with candidate CDSs. Some of these have been described previously in other TPA genome annotations as previously mentioned (S10 Table), while the majority showed no significant homology (blastn, tblastn). These putative genes were relatively small (potentially coding for up to 300 amino acids) and many overlapped with already annotated regions in NC_021490.2. Existence of overlapping genes and short protein-coding CDSs has been experimentally proven in other bacteria before (for instance in *Pseudomonas aeruginosa, E. coli* and *Staphylococcus aureus*) [27-29]. With the growing advances in technologies it becomes clear that small microbial peptides are more abundant than scientists previously thought and that they may play an important role in the bacterial genomes [30]. This is connected to the fact that most of the current bacterial genome annotation programs disallow overlapping CDSs of genes and discard small CDSs below a specific size threshold. Future research should conduct more comprehensive analyses of these transcripts matching with candidate CDSs to better understand their role. Techniques such as mass spectrometry, ribosome profiling, and machine learning predictions will be crucial in these investigations.

Limited transcriptional data from TPA have been reported so far [10, 18]. We hope that this in-depth analyses of TP transcriptome will lay the groundwork for researchers to explore the gene transcription in TPA in our joint effort to elucidate the gene expression of this fastidious and unique pathogen.

## Materials and Methods

### In vitro culture and genome sequencing of TPA Nichols

The TPA strain Nichols sequenced in this study was provided by Steven Norris and Diane Edmondson from the University of Texas Health Science Center at Houston, USA. This strain has been cultured in laboratory animals since 1912 and *in vitro* for several months before being received by our laboratory. It has now been cultured in our lab for over two years according to the protocol designed by Edmondson and colleagues [8] DNA was extracted using the Qiagen DNeasy Blood and Tissue kit (Hilden, Germany) according to the manufacturer’s instructions. The libraries were prepared using the NEBNext Ultra II DNA Library Preparation Kit (New England Biolabs), followed by hybridization with 120-mer RNA baits (SureSelect Target Enrichment System, Agilent Technologies; bait design ELID ID 0616571), as previously described [13]. The enriched libraries were then sequenced on an Illumina Novaseq SP, producing 150 bp paired-end reads (Wellcome Sanger Institute, Cambridge, UK). A total of 11.6 million reads were generated. To identify SNPs and structural variations compared to the TPA Nichols genomic re-sequenced reference from 2012, we used the computational pipeline Breseq (version 0.37.1) [31], optimized for analyzing short-read resequencing data in microbes. To construct the custom reference genome for subsequent analyses, we transferred all annotations from the RefSeq NC_021490.2 (updated as of 03/2023) and incorporated the aforementioned variations and their consequential annotation changes (using Geneious). Locus tag IDs as originally described in the TPA reference genome CP004010.2 [17] were used wherever possible. This reference was used for TPA transcriptome reconstruction and is attached to this study in GenBank format as S1 File. The raw data has been deposited to the European Nucleotide Archive under the project PRJEB49526 (ERP134038) with the Run Accession Number ERR13193639.

### Transcriptome sequencing

To reveal a higher breadth of transcripts expressed in TPA cells, we extracted RNA from TPA cultivated under standard *in vitro* co-culture conditions, as well as incubated in media without the presence of mammalian cells (“axenic culture”). The sample, stored at -80°C with 10% glycerol, was thawed, and one million bacterial cells were inoculated into each culture well. Cultures were grown in TpCM-2 medium under microaerobic conditions at 34°C for 7 days, both in the absence of Sf1Ep cells (axenic culture) and in co-culture with Sf1Ep cells. After 7 days, TPA cells were counted using dark field microscopy. In the co-culture, over 10 million TPA cells were observed with 95% motility, while the axenic culture showed about 500,000 cells with less than 70% motility. RNA was isolated with an RNeasy Mini Kit from Qiagen (Hilden, Germany) according to the manufacturer’s instructions. The concentration and integrity of the RNA were assessed using capillary electrophoresis TapeStation (High Sensitivity RNA ScreenTape Analysis, Agilent, UK).

Multiple different treatments were performed before library preparation to ensure the transcripts would not be compromised. The “co-culture” RNA sample was treated with an NEBNext® Poly(A) mRNA Magnetic Isolation kit (New England Biolabs, UK), and 100 µl of the produced supernatant (bacterial fraction) was used in the subsequent workflow. Ribosomal RNA was depleted using NEBNext® rRNA Depletion Kit (Bacteria) and NEBNext® rRNA Depletion Kit v2 (Human/Mouse/Rat) with RNA Sample Purification Beads (New England Biolabs, UK). An equimolar mixture of depletion solutions from each kit was used during the hybridization step to deplete both eukaryotic and prokaryotic rRNA simultaneously. A second aliquot of “co-culture” RNA was treated only with eukaryotic ribo-depletion probes to preserve potential TPA transcripts. The “axenic” RNA was subjected to prokaryotic ribo-depletion only. Three directional RNA libraries were then prepared: (I)from both ribo- and poly(A)-depleted “co-culture” RNA, (II) from only ribo-depleted “co-culture” RNA, and (III) from ribo-depleted “axenic” RNA (Fig 1). The libraries were prepared using a NEBNext® Ultra™ II Directional RNA Library Prep Kit for Illumina® (New England Biolabs, UK). Each library was deep sequenced on an Illumina NovaSeq to generate 30 Gbp of data per sample (100 million paired-end 150 reads) (Wellcome Sanger Institute, Cambridge, UK). This workflow was inspired by the bacterial-specific RNA-seq previously described for Chlamydia [32].

To reconstruct TPA transcriptome, the RNA-seq data from all three libraries were first merged and mapped to the new custom reference (Bowtie2, version 2.3.5). Aligned reads were then deduplicated (Picard software, version 2.22.2). Only reads with a mapping quality score > 2 were considered further. Merged reads from the RNA sequencing were deposited to the European Nucleotide Archive under project PRJEB49526 (ERP134038) and linked to the files with corresponding reads from the DNA sequencing (Run Accession Number ERR13331577). Genome regions biased by mapping of short reads generated by NGS (e.g., repetitive and paralogous regions) were omitted from the analyses. Only primary aligned reads (excluding reads with multiple alignment) were analysed. Subsequently, 1775 strand-specific coverage plots were generated for all annotated regions of the genome and all intergenic regions larger than 10 bp using a custom Python script. The plots can be accessed at https://tpa-coverage-plots.pam.sanger.ac.uk/ (read coverage details in the S6 Table). Visual inspection of these strand-specific coverage plots allowed for identification of conflicts in annotation as well as for verification of the presence of mRNA corresponding to each annotated regions. Our work-in-progress Nextflow pipeline (TpRNA-seq) is being developed for our future dual RNA-seq studies that are outside the scope of the current study. However, it can already be used to reproduce the TPA transcriptome analyses and strand-specific coverage plots (https://github.com/sanger-pathogens/TpRNAseq). Finally, ANNOgesic (version 1.0.22) [33] was used to predict the transcriptional organization based on the genomic sequence, transcriptional profiles and prediction of Rho-independent terminators.

## Acknowledgements

We are grateful to Lorenzo Giacani (University of Washington, Seattle, US), for critical review of the manuscript. We thank Juraj Bosak and David Šmajs (Masaryk University, Czechia), Steven Norris, and Diane Edmondson (University of Texas Health Science Center at Houston), and Austin Haynes (University of Washington, Seattle, US), for their valuable advice while culturing *T. pallidum in vitro*. In addition, we would like to thank Magnus Manske (Wellcome Sanger institute, UK) for helping to create the web page for the TPA coverage plots.

## Supporting information captions

**S1 Table. SNVs and protein effects found in the genomic sequence of TPA strain Nichols**

**S2 Table. Paralogous regions with ambiguous mapping**

**S3 Table. Annotated region with no evidence of transcription**

**S4 Table. Genes in NC_021490.2 genomic reference coding for hypothetical proteins**

**S5 Table. Potential vaccine candidates**

**S6 Table. Strand-specific transcription tracks details for all annotated regions of the genome and all intergenic regions larger than 10 bp**

**S7 Table. Regions with antisense transcript originating within annotated gene**

**S8 Table. Regions transcribed exclusively in the opposite orientation to the original annotation**

**S9 Table. Regions with transcripts arising from the intergenic region**

**S10 Table. Transcripts with open reading frames corresponding to annotations found in other *Treponema* spp. genomes but absent in NC_021490.2**

**S11 Table. Operons identified in TPA genome**

**S1 File. TPA Nichols annotation reference**

